# The semantic control network mediates the relationship between symbolic numerical order processing and arithmetic performance in children

**DOI:** 10.1101/791012

**Authors:** Gerrit Sommerauer, Karl-Heinz Grass, Roland H. Grabner, Stephan E. Vogel

## Abstract

Behavioral and neuroimaging studies have recently demonstrated that symbolic numerical order processing (i.e., deciding whether numbers are in an increasing/decreasing sequence or not) may engages different cognitive mechanisms and brain regions compared to symbolic numerical magnitude processing (e.g., deciding which of two numerals is larger). Because of this behavioral dissociation, growing interest has emerged to better understand the neurocognitive mechanisms of symbolic numerical order processing and their relationship to individual differences in arithmetic performance. In the present functional imaging work, we further investigated this link in a group of thirty children (7.2-10.25 years) from elementary school, who completed a symbolic numerical order verification (are the numbers going up? e.g., 1-2-3), a symbolic numerical magnitude comparison task (which is the larger number? e.g., 5-7), as well as an arithmetic fluency test outside the scanner. Behavioral results demonstrated the unique role of numerical order to predict children’s arithmetic skills and confirmed its mediating power to explain the association between numerical magnitude and arithmetic performance. Imaging results showed a significant association between numerical order and arithmetic in the intersection of the right inferior frontal gyrus and insula, as well as the posterior middle temporal gyrus. An age-dependent change in brain activity was found in the left intraparietal sulcus. These findings solidify the developmental importance of symbolic numerical order processing in children and provides new evidence that the semantic control network mediates the relationship with arithmetic performance.

**Highlights:**

1. Reaction times of numerical order are a unique predictor of arithmetic (73)
2. Numerical order mediates the relationship of numerical magnitude with arithmetic (83)
3. Brain activation of numerical order processing changes with age in the left IPS (82)
4. The semantic control network mediates the relationship with arithmetic (79)

## 1. Introduction

Symbolic numerical abilities build a crucial foundation for the development of arithmetic competences (Schneider et al., 2017). While past research has mainly focused on symbolic numerical magnitude processing (i.e., the knowledge that a numeral refers to a number of objects, e.g., the digit 7 might refer to 7 cookies; (Sokolowski et al., 2017; Vogel & Ansari, 2012), another important symbolic dimension has received considerably less attention—the dimension of numerical order (Lyons et al., 2016). Symbolic numerical order can be defined as the knowledge that number symbols contain information about their relative rank or position within a numerical sequence (Lyons et al., 2016; Nieder, 2005). For instance, knowledge about the numerical order of the digit 7 allows for an efficient judgement that this digit 7 comes right after 6, but before 8. Although the conceptual distinction of numerical magnitude and numerical order is known for a long time, only recently behavioural and neuroimaging studies demonstrated that these concepts constitute separate cognitive dimensions. In addition, several behavioural studies showed that numerical magnitude and numerical order processing explain unique variance in arithmetic abilities of children (Lyons et al., 2014; Sasanguie & Vos, 2018; Vogel et al., 2014) and adults (Vogel et al., 2017b, 2019). Thus providing, further evidence for a crucial differentiation between the two concepts and the value of numerical order to predict arithmetic abilities. Despite these crucial insights, not much is currently known about the neurocognitive mechanisms that are associated with the development of numerical order processing, how these contrast to the development of numerical magnitude and how the neural correlates mediate the relationship between numerical order and arithmetic in children.

Studies that have investigated numerical order and numerical magnitude processing, have demonstrated distinctive behavioral and neural activation patterns. Symbolic numerical order processing is typically investigated with an ordinal verification task (Goffin & Ansari, 2016; Lyons et al., 2016; Vogel et al., 2017b, 2019). In this task, participants are asked to decide as fast and as accurately as possible whether three presented numerals are displayed in a correct order (e.g., 2-3-4) or not (e.g., 3-2-4). Reaction time measures associated with this ordinal processing show a reverse distance effect: that is numbers with an inter-item distance of one (e.g., 2-3-4) are solved faster compared to trials with an inter-item distance of two (e.g., 2-4-6). Importantly, the behavioral characteristics pattern of the reverse distance effect are significantly different from the well-known canonical distance effect—larger reaction times for small distances (e.g., 2-3) compared to large distances (e.g., 2-8)—typically observed during number comparison tasks (Goffin & Ansari, 2016; Turconi et al., 2006). This behavioral difference has been taken as evidence that numerical order processing engages different cognitive representations (i.e., knowledge about the numerical order and knowledge about numerical magnitudes).

In line with these findings are neuroimaging studies with adults that have shown that numerical order processing also engages different brain mechanisms compared to numerical magnitude processing (Franklin & Jonides, 2009; Lyons & Beilock, 2013; Turconi & Seron, 2002; Turconi et al., 2004). For instance, Franklin and Jonides (2009) were among the first to observe that the neural correlates of numerical order and numerical magnitude differ within a core region of numerical information processing—the intraparietal sulcus (IPS), which is typically associated with numerical magnitude representations (Vogel et al., 2015, 2017a). Only one study has directly contrasted developmental differences in the neural correlates of numerical order and numerical magnitude processing (Matejko et al., 2018). Using a number comparison and a numerical order verification task, the authors contrasted the brain response of children and adults in fMRI. Results of this developmental neuroimaging study showed that adults engaged the left inferior parietal cortex during numerical order processing, while children exhibit brain activation in the right lateral orbital and inferior frontal gyri (IFG) for numerical order and numerical magnitude processing. In line with the results from adults, the authors interpreted these age-dependent differences as evidence for a developmental differentiation of numerical order and numerical magnitude processing in the inferior parietal cortex—especially in the IPS, which was argued to be tuned for numerical order processing with increasing age. Although a significant developmental differentiation was observed in this study, the coarse comparison between children and adults does not allow for a fine-grained developmental differentiation between numerical order and numerical magnitude representations in children (Karmiloff-Smith, 2010). Studies that investigate developmental changes within critical age ranges are needed in order to better understand developmental trajectories of numerical order processing.

There is evidence that numerical order processing undergoes an important change within the first years of formal education. Recent findings from behavioral studies have shown (a) that the associative strengths between numerical order and arithmetic and between numerical magnitude and arithmetic express different developmental trajectories in children attending 1^st^ and 6^th^ grade—the strength between numerical order and arithmetic increases during the first years of formal education, while the strength between numerical magnitude and arithmetic decreases (Lyons et al., 2014; Vogel et al., 2014) and (b) that numerical order processing mediates the well-established association between symbolic numerical magnitude processing and arithmetic in children, and that this mediation effect emerges during the first years of formal education (Sasanguie & Vos, 2018). This evidence suggests a developmental change in the representation of numerical order and its association with arithmetic abilities in the first years of formal education. Thus, neuroscientific studies on developmental trajectories of numerical order processing should focus on this specific age range.

To the best of our knowledge only two fMRI studies exist that have investigated the neural correlates of numerical order processing and its association with arithmetic abilities in children. Both studies contrasted the brain activation of children with and without mathematical learning difficulties (MD, (Kaufmann et al., 2009; McCaskey et al., 2018). Kaufmann and colleagues (2009) compared the neural correlates of numerical order processing in 6 typically and 6 atypically developing children (mean age 10.5, SD 2.0) who have been identified with mathematical learning difficulties (MD). Results of this study showed stronger activations in the anterior cingulate gyrus, the right inferior parietal regions (including the IPS) and the supramarginal gyrus (SMG) in children with MD. The authors interpreted this activation increase in children with MD as compensatory mechanisms to be able to perform the task. In a longitudinal study, McCaskey and colleagues (2018) found an age dependent (8-11 years) activation increase in the IFG, the middle frontal gyrus and the left IPS in 23 children with MD (consistent with a compensatory account). However, no significant age dependent changes in brain activation were found in 12 typically developing peers. The increase in brain activation of children with MD might suggest a greater engagement of additional cognitive control mechanisms to compensate for their ability to access relevant numerical information. Besides the small samples, the current evidence is restricted to contrasts between typical and atypical populations, with a focus to identify alterations in children with MD. Information about the normative brain development associated with numerical order processing and its association with arithmetic abilities is still missing.

Although the behavioral evidence indicates a systematic association between numerical order processing and arithmetic, we do not know the cognitive mechanisms that drive this relationship. One possible explanation, which has been recently put forward by Sasanguie and Vos (2018), argues that the association between numerical order processing and arithmetic may be related to similar strategies to retrieve semantic associations from long-term memory. Indeed, several authors have argued that the significant facilitation in reaction times (i.e., reverse distance effect) in numerical order processing may be related to the fast retrieval of consecutive items from long-term memory (Franklin et al., 2009; Lyons & Beilock, 2013; Vogel et al., 2017b, 2019). Since arithmetic relies on similar retrieval strategies, a potential link between numerical order and arithmetic could be explained by these similar mechanisms. A brain network that is associated with the coordination of semantic information retrieval, is the semantic cognitive control network. The semantic control network includes a number of different frontal and parietal brain regions, such as the inferior frontal gyrus (IFG), the posterior middle temporal gyrus (pMTG), the IPS, pre-supplementary motor areas and the anterior cingulate cortex. These rather domain-general brain regions have been shown to interact with domain-specific representations to engage in control processing mechanisms that “…are suited to the immediate task or context.” (Ralph et al., 2016, p.8). Brain activations within these regions have been also observed during numerical and arithmetic processing (see meta-analysis from Arsalidou & Taylor, 2011) and have been often equated with working memory (Song & Jiang, 2006) as well as higher cognitive monitoring mechanisms—especially in the context of information manipulation (Christoff & Gabrieli, 2000). If similar cognitive strategies mediate the association between numerical order processing and arithmetic, a significant association between the neural correlates of numerical order processing and arithmetic might be expected in brain regions of the semantic control network. Investigating the normative neurocognitive development of numerical ordinal processing in children of a specific age range, in which the association between numerical order and arithmetic appears to develop, would overcome some of the above discussed shortcomings of previous studies and provide new evidence about the neurocognitive development of numerical order processing and its association to arithmetic abilities.

To this end, we asked a group of healthy young children from 2nd^t^ to 4^th^ grade to complete a numerical order verification, a numerical magnitude comparison task, and a paper-pencil test of arithmetic fluency. Based on the literature, we expected that reaction time measures of numerical order and numerical magnitude would be both significant predictors of arithmetic performance. However, we also expected that numerical order explains unique variance over and above numerical magnitude processing and mediates the relationship between numerical magnitude and arithmetic performance. On the neural level, we aimed to identify the brain regions that mediated the relationship of numerical order processing with arithmetic performance and age. We expected that the association of numerical order and arithmetic should be mediated by regions of the cognitive control brain network described above. Based on the limited information from previous work, we expected to find significant brain-behavior associations in frontal and/or parietal regions-especially in regions of the IFG (Matejko et al., 2018; McCaskey et al., 2018) independent of age effects. We also expected to find significant age related changes in the IPS within this sample of children, as has been shown in the comparison between children and adults (Matejko et al., 2018).

## 2. Methods

### 2.1. Participants

A total of 36 children were recruited from elementary schools. Of this sample, one child had to be excluded because of a discovered brain anomaly and five children had to be excluded because of excessive head motion in the scanner (deviation greater than 3.5 mm from the first volume or a greater than 3 mm jump between subsequent volumes; adopted from Vogel et al., 2015), too many errors in one of the experimental tasks, or missing functional imaging data (e.g., abortion of scanning session). The final data set included 30 healthy children (12 females) - aged between 7.2 and 10.25 years (mean = 108 months (9.01 years), SD = 13.1 months). Most participants were right-handed (five left-handers) and native German speakers with normal or corrected to normal vision. The research project has been approved by the local ethical committee. All participants and their caregivers gave written informed consent. Children were reimbursed with a 40 Euro gift card for a local toy store. Data and analyses scripts of this study are available on the open science framework (OSF): https://osf.io/q6k38/

### 2.2. Tasks and Stimuli

Children performed a computerized numerical order verification task, a non-numerical verification control condition, a computerized numerical magnitude comparison task, and a non-numerical comparison control condition. The numerical order task was adapted from Vogel and colleagues (2017b) and consisted of three single-digit Arabic numerals that were horizontally presented on a computer screen (see Figure 1). Children were asked to indicate as fast and as accurate as possible whether the presented number triplets were arranged in an ascending order (e.g., 3-4-5, 2-3-4) or not (e.g., 3-5-4; 3-4-2). The inter-item numerical distance (i.e., distance 1) was kept constant (e.g., 3-4-5, 2-3-4) across all trials. In the corresponding non-numerical control condition, three symbols (numerals were cut in pieces and rearranged; for a similar approach see Kaufmann et al., 2009) were repeatedly presented on the screen (see also Figure 1). In this task, children were asked to decide whether the symbols were presented in the same colour or not. A simple colour control tasks were chosen to control for general cognitive and perceptive components (e.g., general reaction time, visual process associated with the presentation of symbols) and to isolate cognitive mechanisms that are associated with numerical order processing. To ensure a good comprehension, response selection was kept constant between experimental tasks and their appropriate control tasks, so that conditions only varied in task instructions. Both tasks consisted of 32 trials each and children used their right index finger to indicate when the stimuli were in order/the same colour, and their left index finger when the stimuli were not in order/different colours.

**Figure 1.**
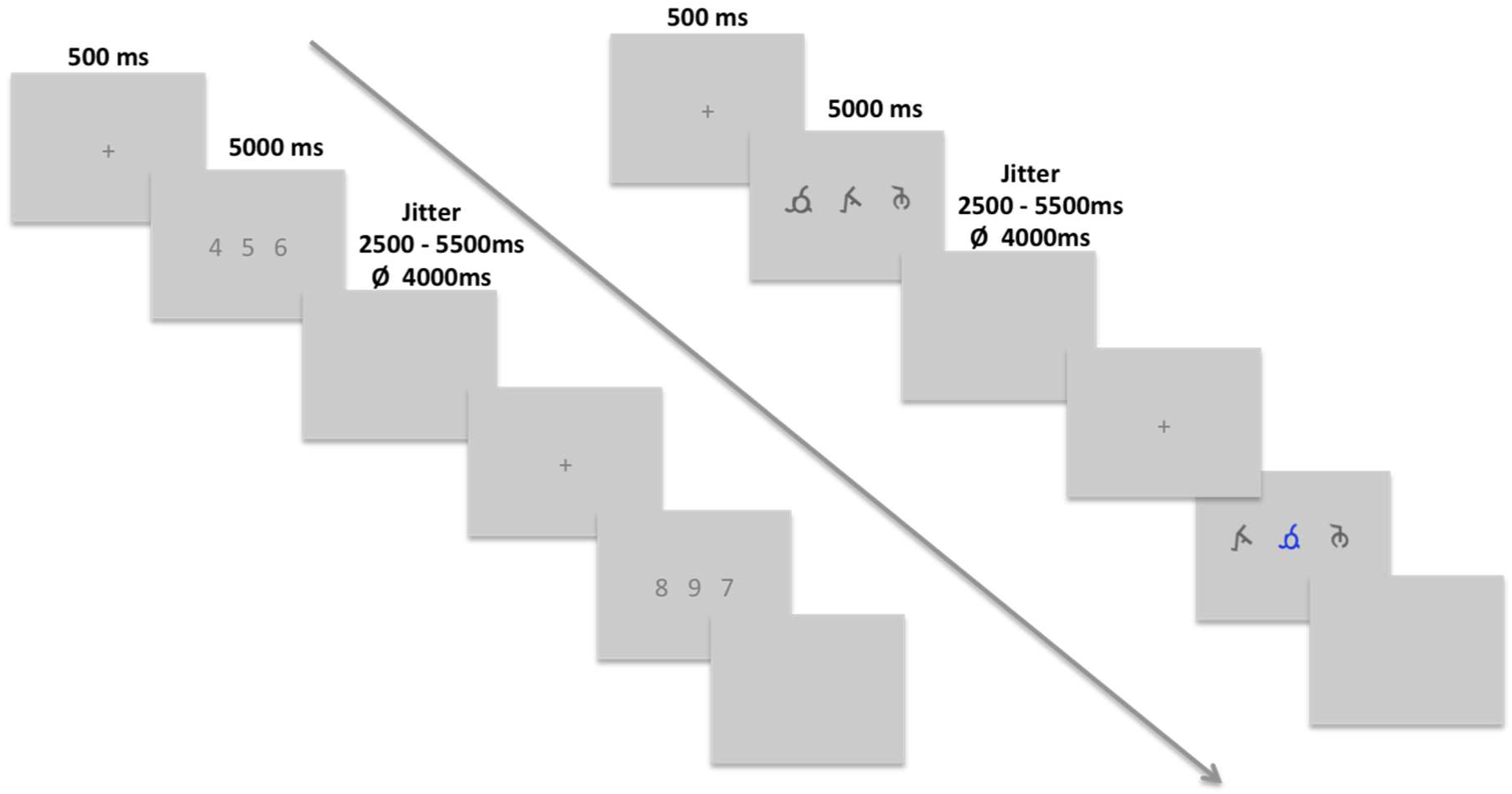
Schematic illustration of a stimulus presentation for the numerical order task on the left side and for the non-numerical task on the right side.

In the numerical magnitude task, two Arabic numerals (e.g., 4 5) were horizontally presented on a computer screen (see Figure 2). Children were asked to decide as fast and as accurately as possible which of the two numerals is numerically larger (i.e., number comparison task; Moyer & Landauer, 1967). Inter-item numerical distances of 1 or 2 (e.g., 3 5; 4 5) were used in this task. In the corresponding non-numerical control condition, two non-numerical symbols (same symbols as in the non-numerical order control condition) were presented on the screen. Children were asked to decide which of the two symbols is printed in blue. Again, a simple colour control task was used to control for general cognitive and perceptive components (e.g., general reaction time, visual process associated with the presentation of symbols) and to isolate cognitive mechanisms that are associated with numerical magnitude processing. Response selection was kept constant between the tasks. Both tasks consisted of 44 trials each and children were instructed to press the right index finger, when the larger number/the blue symbol was on the right side, or, the left index finger when the numerically larger number/the blue symbol was presented on the left side of the screen.

**Figure 2.**
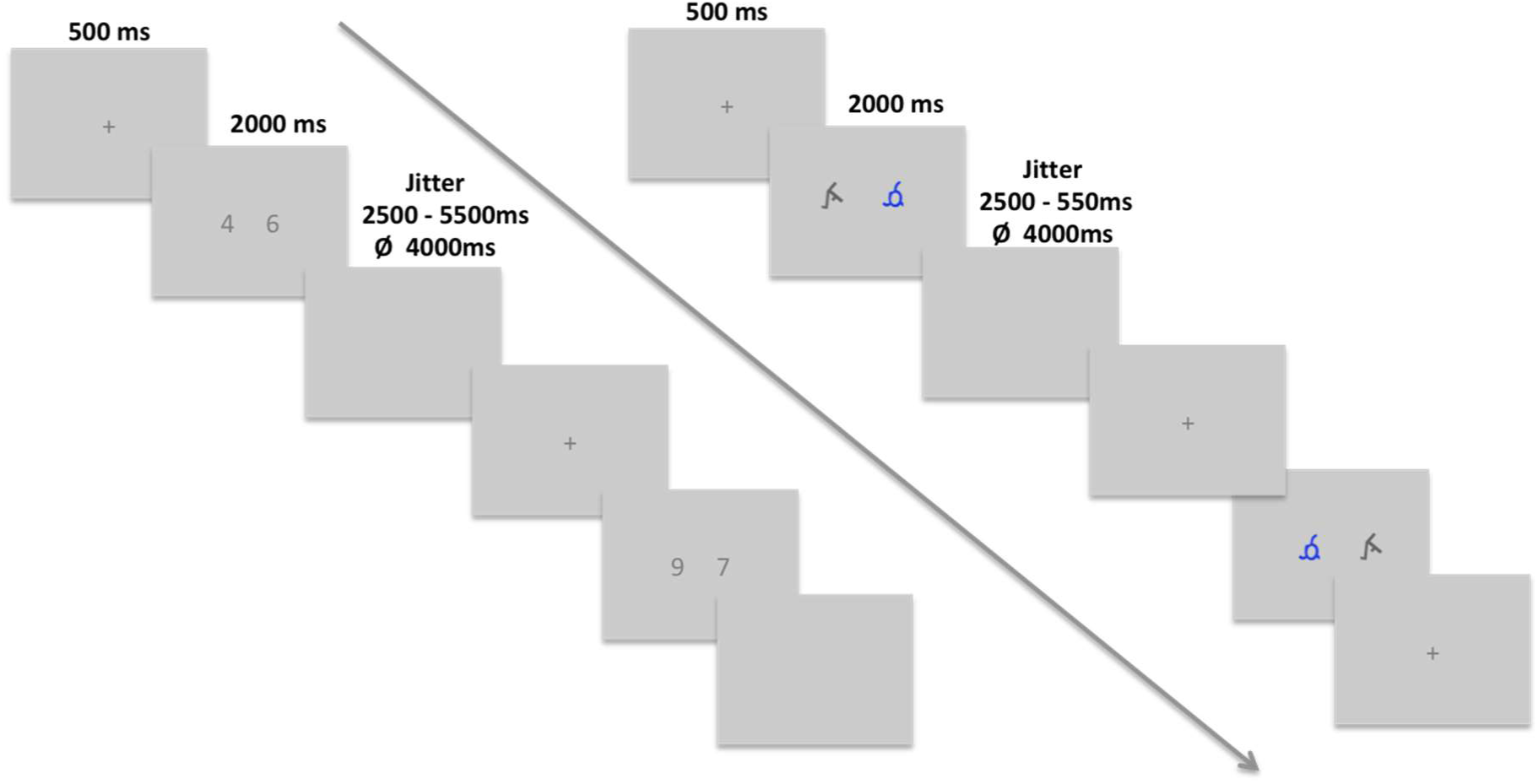
Schematic illustration of a stimulus presentation for the numerical magnitude on the left side and for the non-numerical control task on the right side.

The tasks were presented in pseudo-randomized order using the presentation software PsychoPy (v1.82.01; Peirce, 2008). Participants watched the presentation over a mirror that was placed on top of the channel head coil. Half of the participants (N = 14) started with the numerical order, followed by the non-numerical order control task, whereas the other half of participants (N = 16) completed the numerical magnitude task first, followed by the non-numerical magnitude control task. Each trial (see Figure 1 and 2) started with a fixation cross that appeared for 500 ms on the computer screen. After the fixation cross, the stimuli were presented in the centre of the screen (5000 ms in case of the numerical order task and 2000 ms in case of the numerical magnitude task). The presentation of all stimuli was followed by a blank screen, with a jittered interval of 2500 - 5500 ms (mean of 4000 ms) to oversample the hemodynamic response function (HRF). The duration of each task was about five minutes.

Arithmetic performance was assessed with a paper-pencil test of arithmetic fluency (adapted from Vogel et al., 2017b) outside the scanner. Children completed two pages of addition and two pages of subtraction problems. Per sheet, participants were asked to solve as many problems as possible within 50 seconds. The items of the addition condition consisted of 90 single-digit problems for which the solution was smaller than 10 (e.g., 6 + 1), and of 60 single-digit problems for which the solution was greater than 10 (e.g., 9 + 8). The subtraction condition included 90 items in which the minuend and the subtrahend were single-digit numbers (e.g., 8 - 3), and 60 items in which the minuend was a double-digit number (range 11-19, e.g., 17 - 5) and the subtrahend a single-digit number (range 1 to 9). An arithmetic fluency score (sum of correct answers) was calculated for further analyses.

### 2.3. Experimental Procedure

After a general introduction and a short visiting tour through the facility, children were asked to participate in a short practice session. In this session, all four tasks (numerical order, numerical magnitude and the corresponding control tasks) were practiced on a computer outside the scanner. We also used a crawl tunnel to simulate the scanning bore and to practice lying still. When children felt comfortable to proceed, we carefully positioned them into the scanner and the experimental test session started. The whole fMRI session took about 40 minutes. After the fMRI session, the arithmetic fluency test was administered outside the scanner in a separate and quite testing room.

### 2.4. fMRI data acquisition

Structural and functional imaging data were acquired with a 3-Tesla Siemens Skyra whole-body MRI scanner at the MRI-LAB, University Graz (https://psychologie.uni-graz.at/en/mri-lab/). Changes in brain metabolism associated with neural activity were collected with a 32-channel head coil, using a blood oxygen level dependent (BOLD) sensitive T2* weighted multiband echo planar (EPI) sequence. Images were acquired in an ascending-interleaved order covering the whole brain with 38 slices per volume (2mm thickness, 64 × 64 matrix (voxel resolution = 2.5mm x 2.5mm x 2.5mm), repetition time (TR): 1500ms, echo time (TE): 52ms, flip angle: 78 °). Each functional run consisted of 225 – 236 volumes. High-resolution T1 weighted anatomical images (176 slices in total) were collected with a MPRAGE sequence (1 × 1 x 1 mm, TR: 2300 ms, TE: 4.25 ms, flip angle: 9 °).

### 2.5. Statistical analyses

#### 2.5.1. Behavioural analyses

After calculating descriptive statistics, we performed a set of analyses to investigate the behavioural relationship between reaction time measures of all experimental conditions (numerical order, numerical magnitude and the corresponding non-numerical control conditions) and children’s arithmetic fluency scores. First, we performed Pearson’s and Bayes^1^ correlation analyses to estimate the zero-order correlation amongst the experimental in-scanner tasks, arithmetic fluency scores and age. To further examine which of the above predictors was uniquely related to arithmetic fluency, a partial correlation, a multiple regression analysis and a Bayesian linear regression were calculated. In the partial correlation analysis, we correlated each variable with arithmetic fluency, while controlling for the effect of the remaining variables. In the multiple regression analysis, arithmetic fluency was used as the dependent variable, whereas all other variables (i.e., age and reaction time measures of the experimental conditions) were entered simultaneously into the model (method: ENTER). In the Bayesian regression model, arithmetic fluency was entered as dependent variable and regressed against all other variables. All possible model combinations were first compared to the null-model including age (baseline model). Bayes factor BF_10_ were calculated that indicate how much evidence there is in favour for the best model compared to the null-model. In a second step, Bayes factor BF_12_ were calculated that indicate how much evidence there is in favour for the best model (as revealed in BF_10_) compared to the other models. For all Bayesian analyses we used a flat prior and the JASP package 0.10.0.0 (JASP Team, 2019) to calculate the Bayes factor. For evaluation purposes, we used the classification scheme of Andraszewicz and colleagues (2015), who categorizes a BF_10_ > 3 as moderate evidence, a BF_10_ > 10 as strong evidence, a BF_10_ > 30 as very strong evidence and a BF_10_ > 100 as extreme evidence.

Finally, to investigate whether reaction times of numerical order are a significant mediator between numerical magnitude processing (predictor) and arithmetic fluency (outcome), we performed a mediation analysis based on the PROCESS macro in SPSS (Hayes, 2013) using the software package JAMOVI (The jamovi project, 2019). Mediation analysis investigates whether an *indirect effect* (written as **ab**) of the mediator(s) accounts for some portion of the *total effect* (written as **c**) observed between the predictor and the outcome variable. The unmediated *direct effect* of the original relationship between predictor and outcome variable is denoted as **c’**. When **ab** (i.e., *indirect effect*) is significant but not **c’**, this is interpreted as *full mediation*. However, when both **ab** and **c’** remain significant, the effect is called a *partial mediation*. In our first mediation model, we defined numerical magnitude as the predictor and arithmetic fluency scores as the outcome. Numerical order and non-numerical order (control condition) were defined as mediator variables. In our second model, we tested the opposite mediation model with numerical order as predictor, numerical magnitude and non-numerical magnitude (control condition) as mediators and arithmetic fluency scores as outcome variable (for a similar approach see Sasanguie, Lyons, Smedt, & Reynvoet, 2017). For both mediation analyses we used the bootstrapping method to assess the significance of the mediation model (Hayes, 2013; Preacher & Hayes, 2008). Bootstrapped confidence intervals (CIs) are constructed for *indirect effects* (**ab**) by creating resamples from the original dataset (5000 iterations in the current study). Conclusions about the significance of a results are based on the estimated confidence intervals. When the confidence interval includes 0, the effect is considered to be non-significant, when the confidence interval does not include 0, the effect is considered to be significant (indicators of classical hypothesis testing are nevertheless included). The bootstrapping method is a robust approach to adjust for violations of normality and is recommended for small sample sizes (Preacher & Hayes, 2008).

#### 2.5.2. Imaging analyses

Functional and structural imaging data were pre-processed and analysed with the software package Brain Voyager QX 2.8.4 (Brain Innovation, Maastricht, The Netherlands; (Goebel et al., 2006)) and with the Matlab-based toolbox Neuroelf (v10; http://neuroelf.net/) to calculate whole-brain regression analysis. All functional runs were pre-processed using a head motion (trilinear/sinc interpolation), a slice time correction, a mean intensity signal adjustment and a temporal filtering with a High-Pass filter (GLM-Fourier, 2 sine/cosine cycles). Furthermore, individual functional data sets were aligned to their corresponding anatomical images. After this co-registration, all functional and anatomical images were manually transformed into Talairach space (Talairach & Tournoux, 1988). Finally, all functional data sets were spatially smoothed using a kernel of 6mm (FWHM). A random effects (RFX) general linear model (GLM) was administered to model the functional events and to perform statistical analyses. The length of each functional event (onset-offset) was modelled individually as a function of reaction time (i.e., variable epoch model). This procedure is known to be suited to detect time-varying signals in event-related fMRI studies (Grinband et al., 2008; Yarkoni, Barch et al., 2009). The two experimental conditions and the two non-numerical control conditions were entered as predictors of interest, incorrect trials and motion correction data were modelled as separate regressors of no interest.

The second analysis examined brain-behaviour relationships of numerical order and numerical magnitude processing with arithmetic fluency and age. The first regression model included arithmetic fluency scores and age as predictors for the contrast “numerical_order_ > non-numerical_control_”, the second model included arithmetic fluency scores and age as predictors for the contrast “numerical_magnitude_ > non-numerical_control_”. Using these regression models, whole-brain analyses were calculated to investigate the associations of numerical order and numerical magnitude processing with age (while controlling for individual differences in arithmetic fluency) and the associations of numerical order and numerical magnitude with arithmetic fluency (while controlling for effects of age). For all brain activations, we used an initial uncorrected threshold of p < 0.005 that was subsequently corrected for multiple comparisons using cluster size thresholding (Forman et al., 1995; Goebel et al., 2006). In this method, the uncorrected maps are submitted to correction criteria based on estimates of the map’s spatial pre-processing smoothness and on an iterative correction procedure (Monte Carlo simulation) that estimates cluster-level false-positives rates. After 1000 iterations, the minimum cluster-size that yields a false–positive rate (α) of 0.05 is used to threshold and to correct the statistical maps. Only activation cluster whose size met or exceeded the cluster threshold were allowed to remain on the statistical maps.

## 3. Results

### 3.1. Behavioural results

Descriptive statistics for reaction times and error rates of the scanner tasks are depicted in Table 1 and Figure 3. Calculated pairwise t-test comparisons (corrected for multiple comparison) showed higher error rates for the numerical order condition (p_tukey_ < 0.05; compared to all other conditions), as well as differences in reaction times across all conditions (p_tukey_ < 0.05). In the arithmetic fluency test, children solved between 13 and 133 calculations (mean = 64.3, SD = 28.8).

**Table 1.** Means, standard deviations and confidence intervals (95%) for reaction time and accuracy data.

| Task | Reaction Times (ms) | Confidence Interval (95%) | Accuracy (%) | Confidence Interval (95%) |
| --- | --- | --- | --- | --- |
| Numerical Order | 2349 (534) | 2149-2548 | 95.10 (5.10) | 93.20-97.00 |
| Non-numerical Order | 1526 (306) | 1412-1640 | 97.80 (3.50) | 96.50-99.10 |
| Numerical Magnitude | 1287 (206) | 1210-1364 | 97.70 (3.01) | 96.50-98.80 |
| Non-numerical Magnitude | 998 (170) | 934-1061 | 98.80 (3.41) | 97.50-100.00 |

Note. Stand deviations are provided in parentheses; ms = milliseconds.

**Figure 3.**
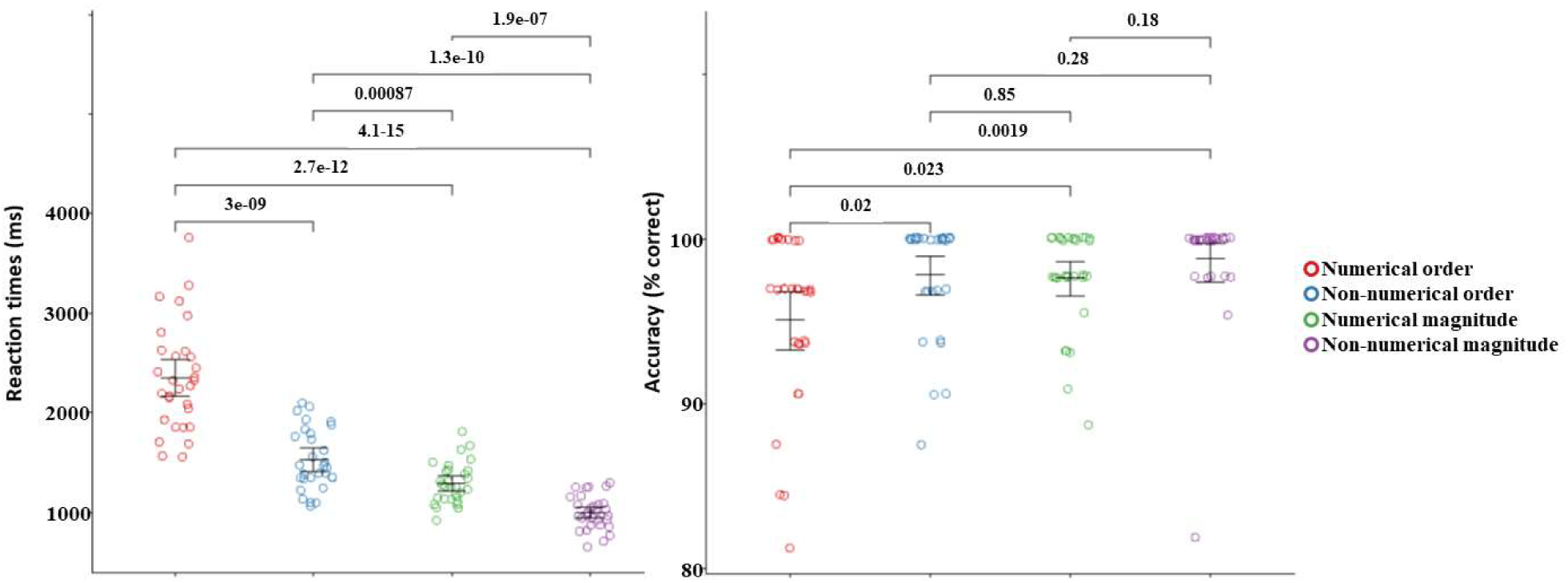
The left illustration depicts the distribution characteristics of reaction time data, the right depicts the distribution characteristics of accuracy data. Individuals’ data are visualized as dots, black bars indicate mean values and 95% confidence intervals; ^***^ p < .001, ^**^ p < .01, ^*^ p < .05; precise p-values for t-tests between conditions are provided on top as suggested by (Amrhein et al., 2019).

#### 3.1.1. Relationship with Arithmetic Fluency

To test the behavioural relationship of numerical order and numerical magnitude with children’s arithmetic performance, we first conducted Pearson’s correlation analyses. Results of these analyses (see Table 2) showed negative correlations between reaction times of numerical order and arithmetic test scores, as well as between numerical magnitude and arithmetic test scores. The better children’s performance in these tasks, the more arithmetic problems were solved. The Bayes factors (see also Table 2) indicated extreme evidence for an association between numerical order processing and arithmetic and very strong evidence for an association of numerical magnitude processing and arithmetic.

**Table 2.** Correlation and partial correlation matrix between all conditions of interest.

|  | 1 | 2 | 3 | 4 | 5 | 6 |
| --- | --- | --- | --- | --- | --- | --- |
| 1 Arithmetic fluency | --- | 14005.718 | 113.245 | 64.540 | 5.538 | 457.299 |
| 2 Numerical order | **-.754** (p < .001)<br>(*-.541\|-.876*) | --- | 146.35 | 7.202e +8 | 243.34 | 16.62 |
| 3 Non-numerical order | **-.614** (p < .001)<br>(*-.326\|-.798*) | **.624** (p < .001)<br>(*.804\|.340*) | --- | 62.288 | 9.169 | 21.004 |
| 4 Numerical magnitude | **-.592** (p < .001)<br>(*-.294\|-.785*) | **.901** (p < .001)<br>(*.952\|.801*) |  | --- | 2230.483 | 1.108 |
| 5 Non-numerical magnitude | **-.466** (p = .009)<br>(*-.127\|-.707*) | **.642** (p < .001)<br>(*.367\|.814*) | **.496** (p = .005)<br>(*.726\|.165*) | **.710** (p < .001)<br>(*.852\|.470*) | --- | 0.748 |
| 6 Age | **.664** (p < .001)<br>(*.826\|.398*) | **-.528** (p = .003)<br>(*-.207\|-.746*) | **-.540** (p = .002)<br>(*-.223\|-.754*) | **-.337** (p = .068)<br>(*.026\|-.622*) | **-.295** (p = .114)<br>(*.073\|-.592*) | --- |

Note. Lower triangle shows correlation values, r-values printed in Bold; p-values printed in parentheses; 95% confidence intervals depicted in Italic. Upper triangle shows Bayes Factors in favour of the alternative hypothesis (BF<sub>10</sub>).

Importantly, the relationship between numerical order and arithmetic performance remained significant when controlling for age, and for reaction times of the numerical magnitude task as well as the two non-numerical control tasks using a partial correlation *and a multiple regression analysis* (see Table 3).

**Table 3.** Arithmetic fluency regressed on all four task variables and age in month.

| Predictor | B | 95% CI<br>(lower\|upper) | SE | t | p-value | r<sub>partial</sub> |
| --- | --- | --- | --- | --- | --- | --- |
| Numerical Order | -0.042 | -0.078\|-0.007 | 0.017 | -2.503 | .020* | -0.455* |
| Non-numerical Order | -0.014 | -0.045\|0.016 | 0.015 | -0.967 | .343 | -0.194 |
| Numerical Magnitude | 0.045 | -0.045\|0.136 | 0.044 | 1.032 | .312 | 0.206 |
| Non-numerical Magnitude | -0.006 | -0.063\|0.051 | 0.028 | -0.219 | .829 | -0.045 |
| Age in months | 0.575 | -0.133\|1.284 | 0.343 | 1.676 | .107 | 0.324 |
| Constant | 72.04 | -30.847\|174.934 | 49.852 | 1.445 | .161 |  |

Note. \* p < .05. Adjusted R<sup>2</sup> is 0.624. Rightmost table column “r<sub>partial</sub>” demonstrates the correlation between one of the predictors and arithmetic fluency scores while controlling for shared variance with all other predictors.

The Bayesian regression analysis (see Table 4) provided additional evidence for this finding. The results showed extreme evidence in favour of a model that contains numerical order compared to the null model (including age) and strong evidence in favour of a model that contains numerical magnitude compared to the null model (including age). A direct comparison between these models showed that the model containing numerical order is 11.421 times more likely compared to the model containing numerical magnitude, and that the parsimonious model containing only numerical order is 3.286 more likely compared to the second best model that contains numerical order + numerical magnitude. Overall, these findings demonstrate that numerical order is a significant predictor that uniquely explains variance of arithmetic test scores over and above the other variables, especially numerical magnitude.

**Table 4.** Results of Bayesian regression model comparison.

| Models | P(M) | P(M\|data) | BF<sub>10</sub> | BF<sub>12</sub> |
| --- | --- | --- | --- | --- |
| Null Model (incl. age in months) | .200 | .014 | 1.000 | -- |
| Numerical order | .050 | .456 | 130.903 | 1.000 |
| Numerical order+Numerical magnitude | .033 | .093 | 39.833 | 3.286 |
| Numerical order+Non-numerical order | .033 | .090 | 38.850 | 3.369 |
| Numerical order+Non-numerical magnitude | .033 | .068 | 29.357 | 4.459 |
| Numerical order+Non-numerical order+Numerical magnitude | .050 | .057 | 16.379 | 7.992 |
| Numerical magnitude | .050 | .040 | 11.462 | 11.421 |

Note. All models include age in months. BF<sub>10</sub> shows the Bayes Factor in evidence for the corresponding model compared to the null-model. BF<sub>12</sub> shows how the factor in evidence for the best model (i.e., Numerical order) compared to the corresponding model.

To determine whether numerical order also mediates the association between numerical magnitude processing and arithmetic fluency, a mediation analysis with numerical order and non-numerical order (control variable) as mediator variables was performed. Results of this analysis revealed that the performance of the numerical order task fully mediated the relationship between numerical magnitude and arithmetic fluency (bootstrap point estimate: a1b1 = -.132, SE = .047, CI = -.224 to -.038, p = .005). The mediating effect of non-numerical order was not significant (bootstrap point estimate: a2b2 = -.021, SE = .016, CI = -.052 to .012, p = .203; see also Figure 4). No significant mediation was found for the opposite model, in which numerical magnitude and non-numerical magnitude were entered as the mediators (bootstrap point estimate: a1b1 = .026, SE = .016, CI = - .005 to .057, p = .110; a2b2 = -.003, SE = .005, CI = -.013 to .007, p = .573). The total and the direct effect between the predictor and the outcome variable were both different from 0 (bootstrap point estimate: c = -.041, SE = .007, CI = -.053 to -.028, p < .001; c’ = -.063, SE = .018, CI = -.099 to -.026, p < .001).

**Figure 4.**
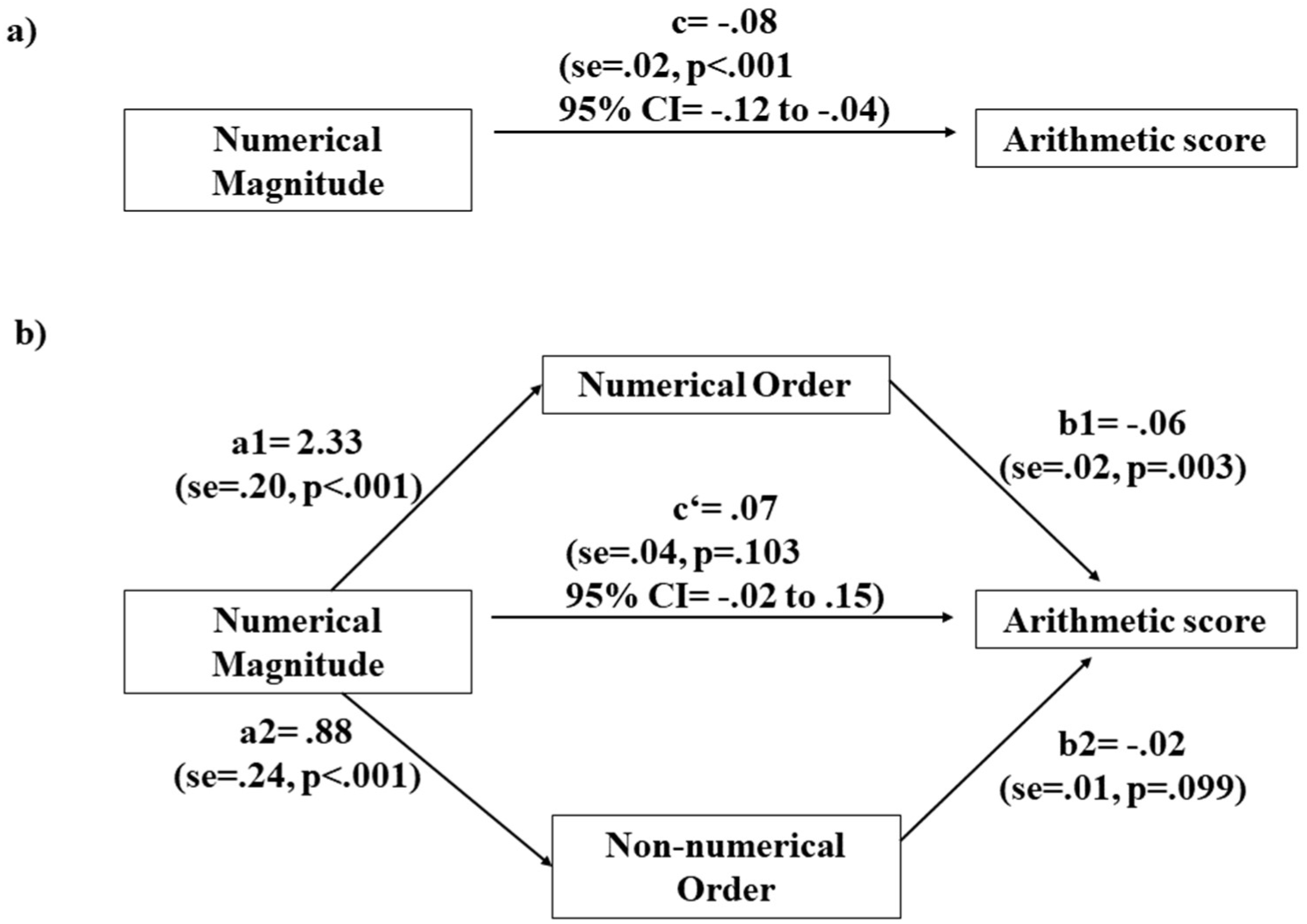
Mediation model. Figure 3a displays the total effect (**c**) of the predictor numerical magnitude performance on the outcome variable arithmetic fluency, after including the mediating variable. Figure 3b demonstrates the mediation effect of the numerical order task (**a1b1**) and the unmediated direct effect (**c’**).

### 3.2. Imaging data

The first analysis aimed to investigate associations between the neural correlates of numerical order, numerical magnitude and arithmetic fluency test scores. Results of this whole-brain regression analysis showed a significant brain-behavior association between numerical order (i.e., numerical_order_> non-numerical_order_) and arithmetic performance in the right posterior middle temporal gyrus (pMTG; Tal_(x,y,z)_: 51, -55, 7; r-value = 0.67; number of voxels = 780) and at the intersection of the right inferior frontal gyrus (opercular part; IFGOp) and the insula (Tal_(x,y,z)_: 42, -1, 16; r-value = 0.70; number of voxels = 1366), while controlling for children’s age (see Figure 6a and 6b for visualization). Children who showed stronger brain activation in response to numerical order processing within these regions achieved higher arithmetic scores in the paper-pencil fluency test. A significant correlation was also found between children’s arithmetic fluency scores and numerical magnitude performance in the left middle frontal gyrus (MFG, Tal(_x,y,z_): -42, 35, 25), after controlling for chronological age.

**Figure 5.**
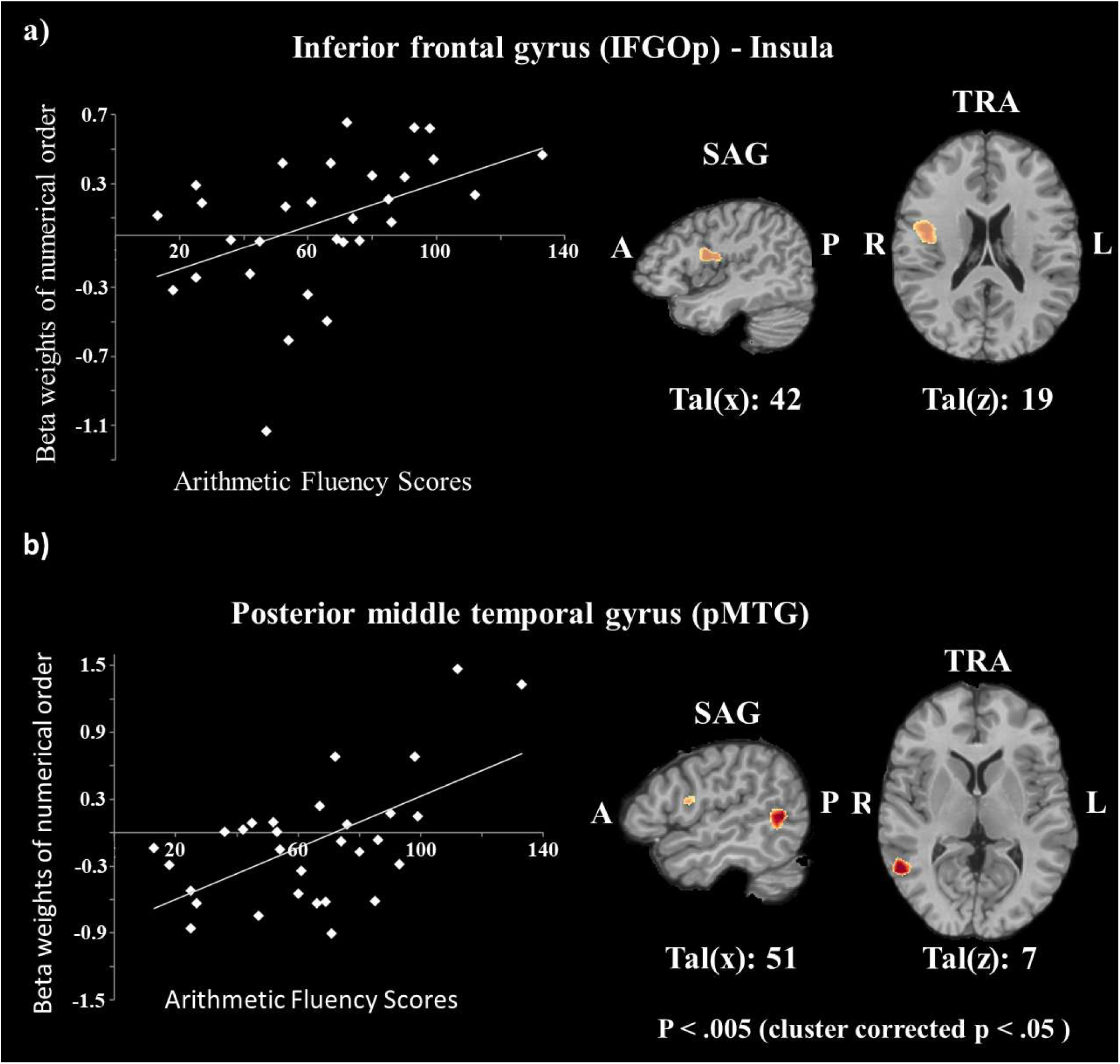
ROI scatterplots depicting the significant correlations between beta weights of the numerical order task and arithmetic fluency raw scores, while controlling for children’s chronological age in a) the IFGOp and the insula and b) the pMTG. Scatter-plots are depicted for illustration purposes. R= right hemisphere, L= left hemisphere, TRA = transversal section. Coordinates are given in Talairach space (Talairach & Tournoux, 1988).

**Figure 6.**
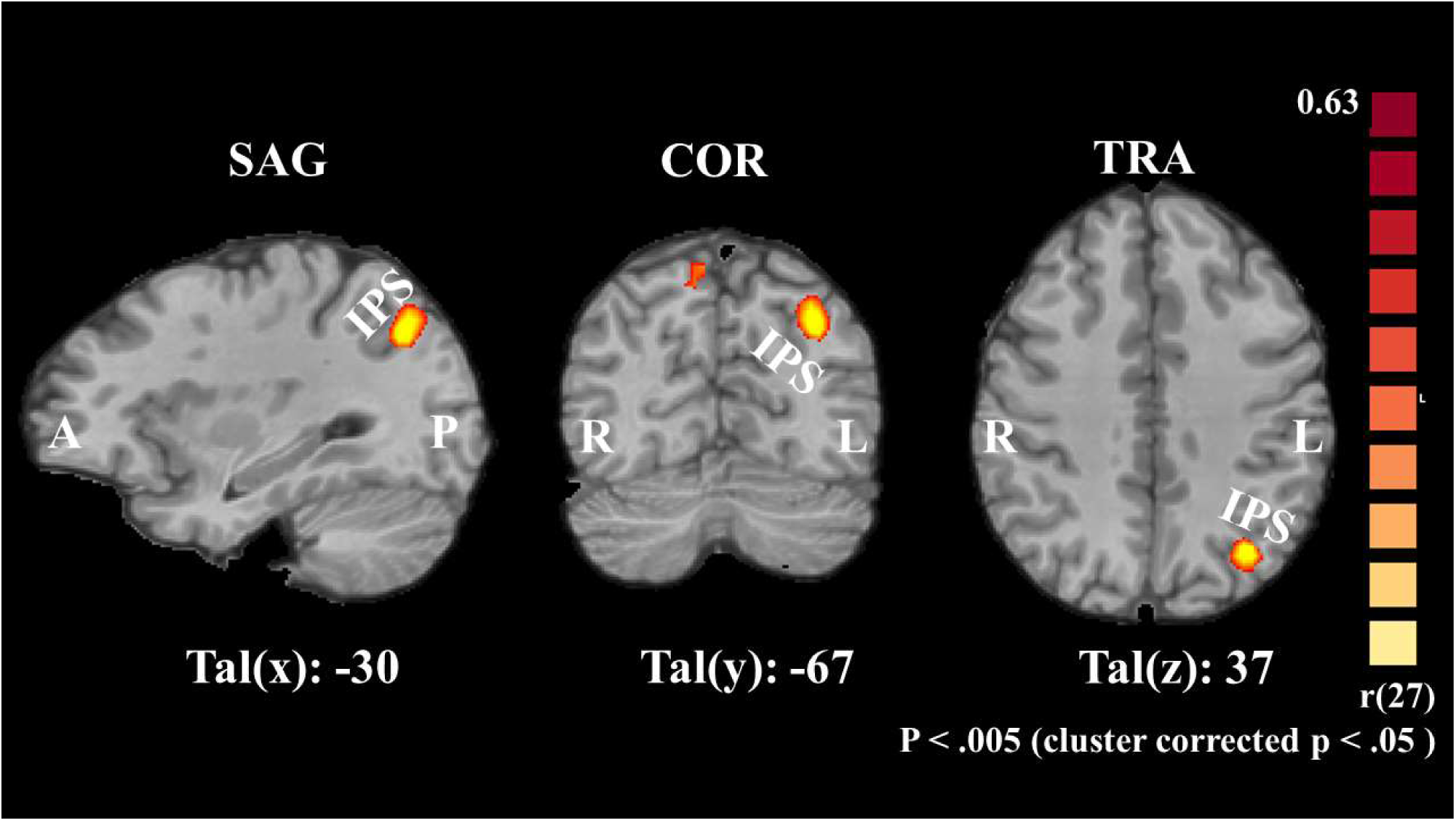
Region of the left IPS that exhibited a significant correlation between numerical order processing and chronological age. The left image shows a sagittal, the middle a coronal and the right a transversal slice of a standardized brain. IPS = intraparietal sulcus; SAG = sagittal; TRA = transversal; COR = coronal; A = anterior; P = posterior; R = right; L = left; Tal = Talairach.

The final whole brain analysis aimed to unravel age dependent changes in the neural correlates associated with numerical order and numerical magnitude processing while controlling for arithmetic abilities. Results of this analysis revealed two brain regions that showed a significant positive association between numerical order processing (i.e., numerical_order_ > non-numerical_control_) and chronological age. The first of these significant clusters (see also Figure 5) was located at the posterior section of the intraparietal sulcus (IPS) of the left hemisphere (Tal_(x,y,z)_: -30, -67, 37; r-value = 0.68; number of voxels = 927), the second cluster was located at the intersection of the precuneus and the superior parietal lobe (SPL) of the right hemisphere (Tal_(x,y,z)_: 6, -55, 67; r-value = 0.67; number of voxels = 823). In contrast, no significant association between numerical magnitude (i.e., numerical_magnitude_ > non-numerical_control_) processing and age was found in this analysis.

Finally, we performed a post-hoc mediation analyses to investigate the specificity of the above identified brain-behavior associations. For instance, it is possible that the significant relationship between numerical order and arithmetic performance in the right IFGOp is mediated by numerical magnitude. Or that the relationship between numerical magnitude and arithmetic in the left MFG is mediated by numerical order. To investigate this possibility, we extracted Beta values from the four brain regions that showed a significant brain-behavior/Age association in the above analyses: (1) the right IFGOp, (2) the right pMTG, (3) the left middle frontal gyrus MFG and (4) the IPS. The extracted Beta values were entered into four separate mediation analysis using the software package JAMOVI (The jamovi project, 2019). The results of these analyses revealed no significant mediation effects. In other words, we found no evidence that numerical magnitude mediated the relationship between numerical order and arithmetic in the right IFGOp (bootstrap point estimate: ab = -.019, SE = .102, CI = -.215 to .183; p = .856) and the rightpMTG (bootstrap point estimate: ab = .005, SE = .038, CI = -.067 to .081; p = .890), that numerical magnitude mediated the relationship between numerical order and age in the left IPS (bootstrap point estimate: ab = -.025, SE = .086, CI = -.184 to .152; p = .775), or that numerical order mediated the relationship between numerical magnitude and arithmetic in the left MFG (bootstrap point estimate: ab = .005, SE = .038, CI = -.067 to .081; p = .890).

## 4. Discussion

The present study is the first to examine the behavioural and neural patterns of numerical order processing and its relationship to arithmetic performance in healthy children attending elementary school. The results of this work demonstrate (a) that behavioural and neural patterns of numerical order processing can be considered significant predictors of arithmetic performance and (b) that the intersection between the right inferior frontal gyrus (opercular part; IFGOp) and insula, as well as the posterior middle temporal gyrus (pMTG) mediate the association between numerical order processing and arithmetic performance.

### 4.1. The behavioural relationship of numerical order processing with arithmetic

Our behavioural analyses showed that the reaction time measures of both numerical tasks (i.e., numerical order and numerical magnitude) correlate with individual performances in arithmetic fluency. This zero-order association is not surprising because it does not control for individual variability of general cognitive mechanisms, such as working memory, which have been shown to be related to arithmetic performance in previous work (e.g., Peng et al., 2016). However, the combined results of the partial correlation and the regression analysis mitigate this confound by demonstrating that children’s ability to recognize the order of numbers relates to arithmetic fluency scores over and above the other experimental conditions (e.g., numerical magnitude and age). Since these analyses investigate the unique contribution of the task, while controlling for other variables and shared variance, a more specific relationship between numerical order processing and arithmetic—over and above numerical magnitude and other domain general components (e.g., age, general reaction time)—is indicated. These behavioural findings are consistent with a growing body of evidence that suggests a unique and reliable association between numerical order judgments and arithmetic skills in children and adults (Goffin & Ansari, 2016; Lyons & Ansari, 2015; Lyons & Beilock, 2011; Lyons et al., 2014; Sasanguie et al., 2017; Sasanguie & Vos, 2018; Vogel et al., 2017b). The results of a mediation analysis further showed that the well-known relationship between numerical magnitude processing and arithmetic is fully mediated by numerical order processing. This finding is strengthened by the inclusion of the non-numerical control condition in the model, which, yet again, controls for domain-general processing components. The result of a full mediation via numerical order processing but not via non-numerical order processing is in line with recent findings from Sasanguie et al. (2017) and Sasanguie and Vos (2018). The first of these studies demonstrated a significant mediation in which numerical order fully explained the relationship between numerical magnitude and arithmetic performance in a group of adults. In the second study, the authors demonstrated a developmental mediating shift in a group of children. More specifically, the authors observed that numerical magnitude mediated the relationship between numerical order and arithmetic abilities in first graders. In second graders, however, numerical order was found to be the significant mediator of the relationship between numerical magnitude and arithmetic. Based on these results, the authors proposed that the association between numerical order processing and arithmetic is driven by similar strategies to retrieve associations from long-term memory: either between numbers (i.e., order) or between problems and solutions (i.e., arithmetic). The results of our study are in line with such an account and show that numerical order mediates the relationship between numerical magnitude and arithmetic in elementary children. Although longitudinal studies are still missing, the emerging picture indicates that numerical order plays a crucial part in the development of arithmetic abilities during the early stages of formal education (see also Lyons et al., 2014; Lyons & Ansari, 2015).

### 4.2. The association between the neural correlates of numerical order processing and arithmetic

The results of the whole-brain regression analysis extend the above discussed evidence by indicating that semantic control mechanisms mediate the neural relationship between numerical order processing and arithmetic fluency. More specifically, the first analysis revealed a significant positive association between numerical order judgments and arithmetic test scores in the intersection of the IFGOp and the insula as well as in pMTG. Children with greater brain activation in both regions had also higher scores in arithmetic fluency while controlling for age. A large body of evidence has observed brain activation in frontal and parietal regions during number tasks (e.g., number comparison) as well as during arithmetic tasks (e.g., subtraction, addition, multiplication; Arsalidou & Taylor, 2011). In their meta-analysis, Arsalidou and Taylor (2011) showed that both conditions (collapsed across addition, subtraction, etc.) engage a larger network of brain regions including the right IFG. Previous neuroimaging work has related IFG activation to visual working memory (Song & Jiang, 2006) as well as to higher cognitive monitoring mechanism—especially in the context of information manipulation (Christoff & Gabrieli, 2000). Because of these characteristics Arsalidou and Taylor (2011) further argued that the IFG is involved in monitoring simple rules that are associated with the manipulation of numerical items. This inference is concordant with the semantic control network (Humphreys et al., 2013). The semantic control network encompasses a variety of brain regions including the IFG and the pMTG, and it is thought that these brain regions interact with domain-specific representations to engage control processing mechanisms that to perform a specific task (Ralph et al., 2016). There is also evidence that the cognitive control network is engaged during the learning of ordinal associations. More specifically, adult participants were asked to learn the ordinal relationship of novel symbols using a transitive inferences task (i.e., from learning x > y and y > z follows the knowledge that x > z; Van Opstal et al., 2009). Results of this imaging study showed a significant increase in the IFG and, at a lower threshold, also in the right pMTG (i.e., contrasting the brain activation of blocks of artificial symbol training during the test phase). This finding indicates that regions of the semantic control network, the IFG and the pMTG, are involved in ordinal sequence learning. In the light of these data, the results of the present work indicate that regions of the semantic control network are involved in mediating the relationship between ordinal knowledge and arithmetic performance. This finding is also consistent with the hypothesis that the association between numerical order processing and arithmetic is related to similar strategies to retrieve semantic associations from long-term memory (Sasanguie & Vos, 2018). If this explanation holds, it also suggests that there might be no direct causal link between ordinal processing and arithmetic, but rather a common third mechanisms that drives the relationship between the two: similar semantic control mechanisms to retrieve relevant numerical information from long-term memory.

### 4.3. The association between the neural correlates of numerical order processing and age

The second whole-brain analysis revealed a significant positive association between numerical order processing and age in the left IPS while controlling for arithmetic performance. In other words, greater brain activation in response to numerical order processing was observed in older kids. While brain activation in the IPS is typically associated with numerical magnitude processing, there is also evidence, primarily from adult studies, that IPS activation is related to the processing of order information. For instance, (Fias et al., 2007) reported similar IPS activation for a number and letter ordinal comparison tasks in a group of healthy adults. Nevertheless, a re-analysis of the same data using multi-voxel pattern analysis (MVPA) indicated a different activation pattern for different numerical and non-numerical ordinal sequences (Zorzi et al., 2011). The finding of an engagement of similar brain regions with different activation pattern was also shown in a study by Franklin and Jonides (2009). The authors investigated the neural correlates associated with numerical magnitude and numerical order processing and reported significant, yet different, activation patterns within the IPS for both conditions. While the authors observed a canonical distance effect for numerical magnitude processing, a reverse distance effect was observed for numerical order processing. These findings indicate an important role of the IPS during numerical ordinal processing. The importance of the IPS for ordinal processing was also recently confirmed by a developmental imaging study (Matejko et al., 2018) in which the authors showed a greater engagement of frontal regions during numerical order processing in children and a stronger activation cluster within the left IPS in adults. Our results replicate and extend this finding. They demonstrate a functional and age-dependent specialisation of the IPS to process ordinal information in the first years of formal education in which knowledge about numerical ordinal processing seems to be established. However, the functional relevance of the IPS seems not to be related to arithmetic performance. Future studies need to further specify the role of numerical order processing in the IPS and its functional relevance.

A potential concern is the fact that in the current task design a numerical ordinal task (i.e., digits) was compared to a colour control task with arbitrary symbols. Thus, it could be argued that the contrast “experimental task > control task” does not solely reflect domain-specific components of ordinal processing (i.e., numerical ordinal processing), but rather components of domain-general ordinal processing (i.e., mechanisms that are not specific to numerical order). In other words, activation found in brain areas such as the IFG may be related to generic aspects of ordinal processing that are not restricted to the processing of numerical relationships. Indeed, there is evidence that processing the order of non-numerical dimensions, such as letters of the alphabet (e.g., D-E-F), demonstrates similar behavioural and neural effects as processing the order of numbers (e.g., 2-3-4; Fulbright et al., 2003; Vogel et al., 2017b). For instance, a similar reverse distance effect can be observed when adults are asked to verify the order of digits and letters of the alphabet (Vogel et al., 2017b). Thus indicating, that similar domain-general components might be associated with numerical and non-numerical ordinal judgments. However, adults appear to make much more errors in the letter ordinal verification task compared to the digit ordinal verification task: in the mentioned study above, an error rate of 27% was found in the letter condition, an error rate of 6% was found in the in the digit condition (Vogel et al., 2017b). This indicates that domain-general aspects of ordinal processing interact with the corresponding knowledge domain to solve the task, and that a clear experimental differentiation between these elements might turn out to be difficult (besides the practical difficulties of implementing a challenging task such as the letter order verification task in an fMRI study with young children). Future work might be able to achieve such a discrimination.

### 4.3. What about the neural correlates of numerical magnitude processing?

The findings of the present work highlight the importance of numerical order processing and its relationship to arithmetic performance. But what about numerical magnitude and its relationship to arithmetic performance? The present work found a significant association between numerical magnitude processing and arithmetic both on the behavioral and the neural level. The behavioral finding is consistent with a large body of evidence that has demonstrated a significant association between numerical magnitude processing and arithmetic (Schneider et al., 2017). The important role of numerical magnitude processing is well established, but the present data suggest that it is not the only important dimension relating to arithmetic abilities. Indeed, it appears that numerical order plays a significant role during the first years of formal education. This general trend is supported by the neuroimaging results. The whole-brain analysis showed that the middle frontal gyrus mediates the relationship between numerical magnitude and arithmetic in elementary school children. Activation in regions of the bilateral middle temporal gyrus are frequently reported in studies with number tasks (Arsalidou & Taylor, 2011). Brain activation in this region has been often associated with attention and working memory (Christoff & Gabrieli, 2000; Curtis & D’Esposito, 2003). In contrast to the IFG, this region is argued to be involved when coordination and cognitive control needs to be applied (Rypma et al., 1999). Arsalidou and Taylor (2011) argued that the activation of the middle frontal gyrus can be related to numbers being held in mind and the operations being applied to them. Especially the dorsolateral region of the prefrontal cortex might be related to monitoring task-relevant information. To the best of our knowledge only one study has investigated the relationship between symbolic numerical processing and arithmetic in healthy children (Bugden et al., 2012). In this work, a significant association between the left IPS and arithmetic abilities was reported. Why do the results of our study not converge with these data? (a) While Bugden et al. (2012) investigated the relationship between the numerical ratio effect and arithmetic, the present work focused on the contrast between numerical magnitude and a corresponding control task. (b) It is entirely conceivable that multiple brain regions facilitate the relationship between numerical processing and arithmetic at different age points. While more domain-general cognitive control mechanisms might be involved in the early stages of development, more domain-specific and automatic mechanisms of number representation might be involved at later stages. The lack of information clearly indicates that we need to do a better job in investigating developmental trajectories across different age groups.

The present work did not reveal age dependent activation changes for numerical magnitude processing. Does this mean that numerical magnitude processing does not change over time? We do not think that this is the case. There is abundant evidence that the neural correlates of numerical magnitude processing changes with age (e.g., Ansari, 2008). For instance, in an fMRI adaptation task we found a numerical ratio dependent interaction with age in the left IPS in response to symbolic numbers in a cross-sectional study including a larger age range of children (Vogel et al., 2015). However, the present work focused on a specific age group to cover changes in numerical order processing during the first years of formal education. Indeed, there is strong evidence that the relationship between numerical order processing and arithmetic changes during this particular time, but not between numerical magnitude and arithmetic (Lyons et al., 2014; Sasanguie & Vos, 2018). For instance, in a large cross-sectional developmental study, Lyons and colleagues (2014) collected a variety of numerical tasks—including numerical order and numerical magnitude—as well as measures of arithmetic performance in children from the 1^st^ to 6^th^ grade. Results of this work demonstrated that numerical order explains increasingly more variance with age, being the best predictor of arithmetic fluency by grade 6. Together with our data, this may indicate that developmental changes in the explored age range are more pronounced for numerical order processing than for numerical magnitude processing—which may be established in years prior to formal education. The different association of numerical order processing and numerical magnitude processing with arithmetic in the present work provides further evidence for the involvement of distinct cognitive mechanisms in these two tasks. The precise dynamic need to be further investigated to better understand the foundational role of symbolic numerical knowledge during the first years of formal education.

### 4.4. Conclusion

In conclusion, the current work showed that behavioural performance of both tasks—numerical order and numerical magnitude—was significantly correlated with arithmetic fluency. Consistent with previous findings, we further observed that numerical order predicted unique variance in arithmetic performance and mediated the association between numerical magnitude and arithmetic performance. Furthermore, our brain imaging data suggest that the semantic control networks is involved in mediating the relationship between numerical order processing and arithmetic. This interpretation is consistent with the notion that similar cognitive strategies drive the observed correlations between numerical order processing and arithmetic performance.

## Acknowledgements

The authors thank Dennis Wambacher for supporting the technical realization of the experiment and his assistance in the data acquisition and Jakob Kelz for supporting the recruitment of participants. fMRI data collection was supported by Dr. Karl Koschutnig and Thomas Zussner, MSc. We like to thank them both very much. Finally, we thank Prof. Daniel Ansari and Prof. Ian Lyons for the constructive comments. Finally, we would like to thank all the families and children who supported this research.

## Footnotes

1 The Bayes factor is a ratio that quantifies how well the data fit under the alternative-hypothesis compared to the null-hypothesis (H_10_) or vice versa (H_01_; Jarosz & Wiley, 2014). As the value of the Bayes factor increases, the evidence in support of the alternative hypothesis, or the null-hypotheses, increases (for further information about the Bayes factor see (Jarosz & Wiley, 2014; Masson, 2011; Wagenmakers, 2007).

